# IgM plays a prominent role in naturally acquired immunity against *Plasmodium falciparum* gametocytes

**DOI:** 10.1101/2024.06.30.601434

**Authors:** Jo-Anne Chan, Ashley Lisboa-Pinto, Shirley Lu, Alexander Harris, Matthew WA Dixon, Adam Thomas, Damien R Drew, Niva Jayakrishnan, Katrina Larcher, Mohammad Naghizadeh, D Herbert Opi, Linda Reiling, Michael Theisen, Kiprotich Chelimo, Maria Ome-Kaius, Daisy Mantila, Moses Laman, Leanne J Robinson, Ivo Mueller, Christopher L King, Arlene Dent, James W Kazura, James G Beeson

## Abstract

The development of transmission-blocking vaccines against *Plasmodium falciparum* malaria could facilitate malaria elimination. However, limitations in the knowledge of the human immune responses against *P. falciparum* transmission stages, known as gametocytes, represent a critical roadblock to vaccine development. We evaluated human antibodies acquired through natural malaria exposure to whole gametocytes and recombinant antigens expressed by transmission stages, including the major transmission-blocking vaccine candidates Pfs230 and Pfs48/45 and other transmission stages, Pf38, Pf12 and Pf41. Among individuals residing in Kenya and Papua New Guinea, we found substantial antibody responses to whole gametocytes and to all recombinant transmission stage antigens with high levels of IgG, IgG subclasses and IgM. Complement fixation by antibodies to gametocytes is key for effective transmission-blocking activity. We found that purified IgM was substantially more potent than IgG at mediating complement fixation and activation. Higher antibody levels were generally observed in individuals positive for *P. falciparum* infection, including gametocyte positive individuals, and these antibodies generally increased with age. Our findings reveal that IgM is a prominent feature of antibody responses to gametocytes and that antibodies target multiple antigens. The further demonstration that IgM has high functional activity against gametocytes suggests IgM plays an important role in immunity to transmission stages. Our data provide new insights to inform the development of potent transmission-blocking vaccines.

## INTRODUCTION

Malaria caused by *P. falciparum* is a substantial global public health burden, with over 240 million cases and half a million deaths annually^1^. The decline in malaria incidence has plateaued since 2015^1^. The licensed malaria vaccine, RTS,S, targets the sporozoite stage of *P. falciparum* (mosquito to human) but not human to mosquito transmission stages, and only confers modest efficacy in young children and infants (55% over 1 year, 36% over 4 years)^2^. It is also not currently recommended for use in older children and adults who are major drivers of malaria transmission^3^. Thus, strategies to develop next generation vaccines against *P. falciparum* transmission remain a high priority. Developing transmission-blocking vaccines is currently recognised by WHO and global partners as a key goal to achieving and sustaining malaria elimination^1^. Transmission-blocking vaccines would likely be used together with pre-erythrocytic or blood stage vaccines that reduce clinical illness, or formulated as a multi-stage vaccine. However, major knowledge gaps in our understanding of the human immune response to transmission stage *P. falciparum* hamper the advancement of transmission-blocking vaccines^4^.

The transmissible sexual stages of *P. falciparum*, known as gametocytes, are distinct parasite forms that circulate in the blood and can be taken up by mosquitoes during a blood meal to establish infection in the mosquito vector. Naturally acquired antibodies against gametocyte surface antigens develop in malaria-exposed individuals (reviewed in ^5^). These antibodies have the potential to reduce malaria transmission by inhibiting mosquito infection. When a mosquito feeds on an infected person, whole blood is ingested, including antibodies and complement proteins present in plasma which have the potential to inhibit the development of parasites within the mosquito midgut and to reduce malaria transmission (reviewed in ^5,6^). Antigens expressed on the surface of the gametocyte plasma membrane play essential roles for infection and replication of parasite forms within the mosquito. Most gametocyte antigens play a role in gamete fertilization in the mosquito midgut, specifically Pfs230 in male gamete fertility through the formation of exflaggelation centres, and Pfs48/45 in the anchoring of Pfs230 on the surface of gametes. These antigens belong to the 6-cysteine protein family and include the less well characterised antigens Pf38, Pf12 and Pf41 which have been recently shown to be expressed in gametocytes^7,8^, but their function and importance as antibody targets remain unclear. While Pfs230 and Pfs48/45 are targets of acquired immunity, studies have demonstrated that serum depleted of antibodies to these antigens is still capable of inhibiting transmission, suggesting that additional antibody targets are involved. For Pfs230, antibodies to the first domain appear most important for transmission-reducing activity and is the focus of vaccine development^9^. For Pfs48/45, the C-terminal 6-cysteine domain has the immunodominant epitope I that is targeted by transmission-blocking antibodies^10^. Pfs230- and Pfs48/45-based vaccines have entered Phase I clinical trials, which have shown that induced antibodies have substantial transmission-reducing activity by standard membrane feeding assay (SMFA)^10–12^.

Studies have established that antibodies to gametocyte surface antigens are acquired following natural *P. falciparum* infections and higher antibody levels are observed upon repeated exposure^13^. Serum antibodies (IgG) from naturally exposed individuals from the Gambia^14^, Tanzania^15^ and other sites in Africa^16–18^ recognised recombinant Pfs230 and Pfs48/45. Antibody levels were associated with an ability to reduce malaria transmission to mosquitoes using a SMFA in a controlled laboratory setting^5,19^.

Antibodies mediate a variety of effector functions, but few have been studied in the context of transmission-blocking immunity. Complement fixation and activation is known to play an important role in mediating effective transmission-blocking activity by antibodies^20^, which has been demonstrated using human polyclonal antibodies, monoclonal antibodies, and antibodies raised in experimental animals^11,20,21^. However, the drivers of effective complement-fixing antibodies remain unclear. Most studies on transmission-blocking immunity have focused on the role of IgG (reviewed in ^6^), and none have investigated the importance of the IgM response. This is a significant gap in knowledge that may be relevant to understanding transmission-blocking immunity and the development of effective vaccines as IgM can be a potent activator of complement. We have previously identified a key role for IgM activity against asexual blood stages of *P. falciparum*^22^.

In this study, we quantified the importance of gametocyte surface antigens as targets of naturally acquired immunity to malaria, with a specific focus on quantifying the IgM response. We evaluated the hypothesis that IgM plays a role in fixing complement on the surface of gametocytes to mediate effective transmission-blocking activity. We quantified serum antibodies in cohorts of children and adults from malaria endemic regions in Kenya and Papua New Guinea (PNG), to represent diverse epidemiological settings. We measured IgG, IgG subclasses and IgM to *P. falciparum* gametocytes and recombinant gametocyte surface antigens. Further, we purified human IgM and IgG from serum samples from malaria endemic regions and assessed their ability to mediate complement fixation and activation on *P. falciparum* gametocytes.

## RESULTS

### Naturally acquired antibodies from across diverse epidemiologic settings recognise antigens on the surface of gametocytes

To evaluate the natural acquisition of antibodies to gametocytes, we measured IgG, cytophilic IgG subclasses (IgG1 and IgG3) and IgM binding to whole gametocytes. These gametocytes were cultured to maturity (stage V) and treated with saponin to lyse the infected erythrocyte (IE) membrane and parasitophorous vacuole (PV) membrane^23^. We previously established this approach and showed that IgG in serum from Kenyan individuals (n=5) could recognise surface antigens expressed on the plasma membrane of intact gametocytes by flow cytometry^23^. Here, we conducted a detailed investigation in diverse populations using ELISA to quantify IgG and IgM to surface antigens on the plasma membrane of gametocytes. Among samples from Kenyan adults (Kanyawegi; n=94), we found that 87.7% had IgG recognition of whole gametocytes (Fig 1A; antibody positivity is defined as antibody levels greater than the upper 95%CI of the mean responses of malaria-naïve negative controls). We measured IgG subclasses, specifically IgG1 and IgG3 responses because of their ability to activate complement, and found that 97.3% and 100% of individuals had IgG1 and IgG3 antibodies to gametocytes (Fig 1A). This suggests that the cytophilic IgG subclasses directed at gametocytes are acquired following natural malaria infections. Further, 71.2% of adults were positive for IgM to gametocytes (Fig 1A), suggesting a potential role for IgM in transmission-blocking immunity that has not been previously defined; prior studies have focused on IgG. No significant differences in antibody levels to gametocytes were observed between samples that were positive or negative for *P. falciparum* infection (Fig 1A). Samples from a different location in Kenya (Chulaimbo; n=18 children and adults) were also tested; 94.4% of individuals had detectable IgG and 55.6% had detectable IgM responses to gametocytes (Fig 1B). A high proportion of individuals were positive for IgG1 and IgG3 responses (Fig 1B; 88.8% for IgG1, 94.4% for IgG3). Adults had significantly higher levels of IgG and IgG1 compared to children (Fig 1B; *p*=0.04, *p*=0.03, respectively). There was a pattern of higher magnitude of IgG3 and IgM for adults compared to children, but this was not statistically significant; however, the sample size limited statistical power.

**Fig 1.**
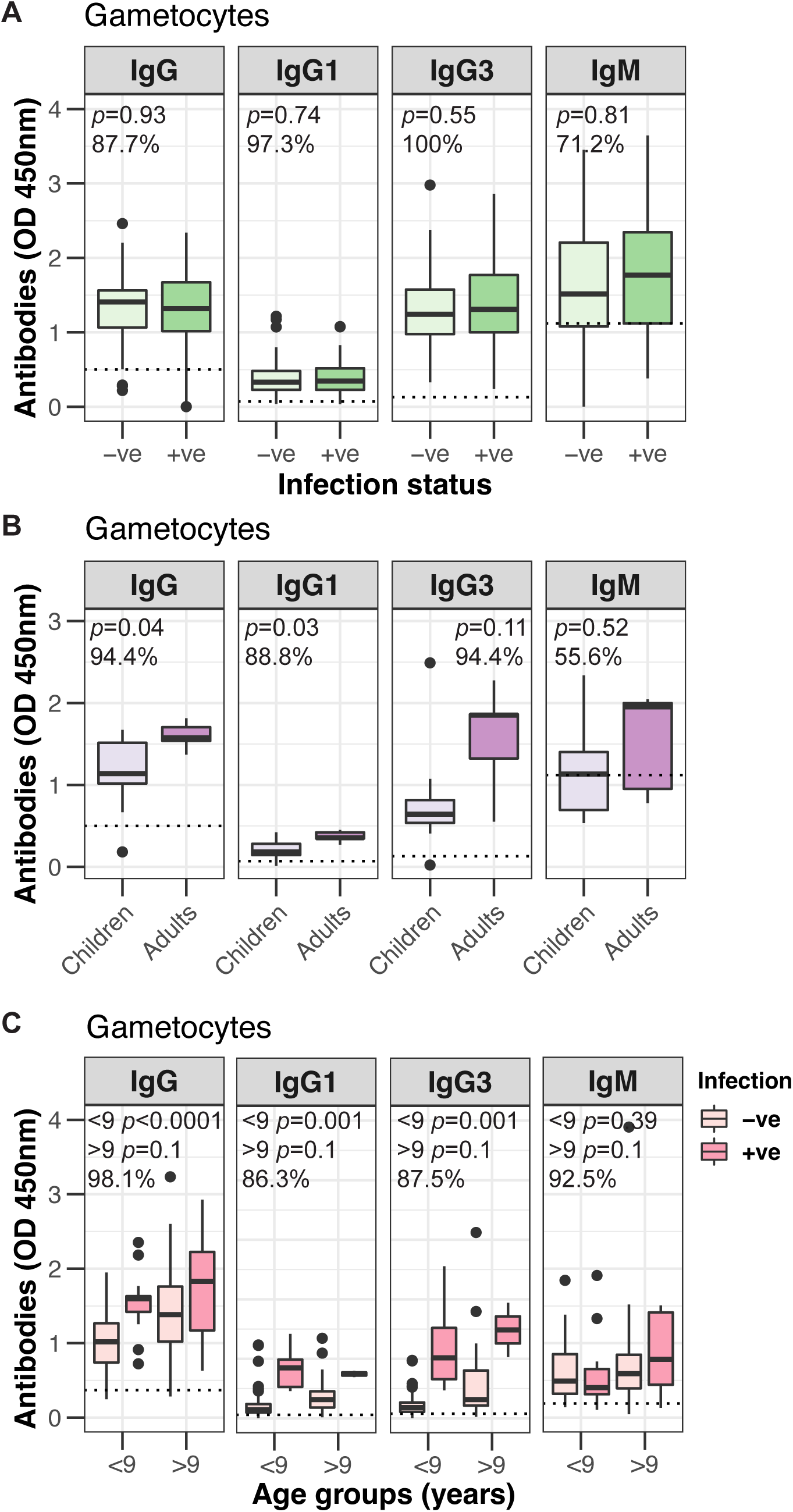
Naturally acquired antibodies to the surface of whole gametocytes. Antibody levels (IgG, IgG subclasses and IgM were measured to the surface of whole gametocytes. (**A**) Samples were from adults (n=94) residing in Kanyawegi, Kenya and are classified according to infection status (+ve, -ve) for each antibody variable. (**B**) Samples were from children (n=13) and adults (n=5) residing in Chulaimbo, Kenya. (**C**) Samples were from children residing in Madang, PNG (n=160 for IgG and IgM, n=80 for IgG subclasses) and divided into 2 age groups (<9 years, n=65 for gametocytes tested for IgG and IgM, n=56 for IgG subclasses; >9 years, n=95 for gametocytes tested for IgG and IgM, n=24 for IgG subclasses) and are classified according to infection status (+ve, -ve) for each antibody variable. Antibody levels were measured by ELISA at optical density of 450nm and tested in duplicate; the dotted line represents the antibody positivity threshold (OD levels greater than the upper 95%CI of the mean responses of malaria-naïve Australian controls); % refers to the percentage of samples above the antibody positivity threshold; *p* values comparing antibody groups (infection status in A and C or children vs adults in B) were calculated using an unpaired Mann Whitney test. Box and whisker plots indicate the first and third quartile for the hinges, median line and lowest and highest values no further than 1.5 interquartile range from the hinges for whisker lines.

To assess antibody responses in a geographically different population, including older children who are a major transmission group in endemic areas^24^, we measured antibody responses in samples from a cohort of children (aged 5-14 years) in PNG^25^. Nearly all children had IgG and IgM reactivity to whole gametocytes (Fig 1C; 98.1% for IgG and 92.5% for IgM). Similarly, 86.3% and 87.5% of children were classified as positive for IgG1 and IgG3 to whole gametocytes (Fig 1C). Higher magnitude of antibodies was observed in older children (>9 years old) for IgG, IgG subclasses and IgM to whole gametocytes compared to younger children (<9 years old) (*p*=0.0003 for IgG, *p*=0.01 for IgG1, *p*=0.01 for IgG3 and *p*=0.18 for IgM), reflecting their greater cumulative exposure to malaria infections. In younger children (<9 years), higher antibody levels for IgG and IgG subclasses were observed for those classified as positive for *P. falciparum* infection compared to those classified as negative (Fig 1C; *p*<0.0001 for IgG and *p*=0.001 for IgG1 and IgG3), but this was not seen for IgM (Fig 1C; *p*=0.39). In older children (>9 years), no difference in antibody levels were observed between those classified as positive or negative for *P. falciparum* infection (Fig 1C; *p*=0.1).

### Naturally acquired antibodies mediate complement fixation on whole gametocytes

Complement plays an important role in mediating transmission-blocking immunity^26–28^ and can be mediated by IgG and IgM. Therefore, we evaluated the functional ability of acquired antibodies to fix complement C1q to whole gametocytes. In serum samples from adults from Kanyawegi, Kenya, all samples were positive for antibody-mediated complement fixation on gametocytes (Fig 2A), though the magnitude of complement-fixing activity varied between samples. No apparent differences were observed between those classified as positive or negative for *P. falciparum* infection (Fig 2A; *p*=0.58). The relationship between antibody variables (IgG1, IgG3 and IgM) and C1q was visualized using a network plot (Fig 2B). IgG1 and IgG3 were clustered closely with C1q (IgG1, r_s_=0.46, *p*<0.0001; IgG3 r_s_=0.71, *p*<0.0001), demonstrating that they are highly correlated, while IgM and C1q were located further apart (r_s_=0.21, *p*=0.08).

**Fig 2.**
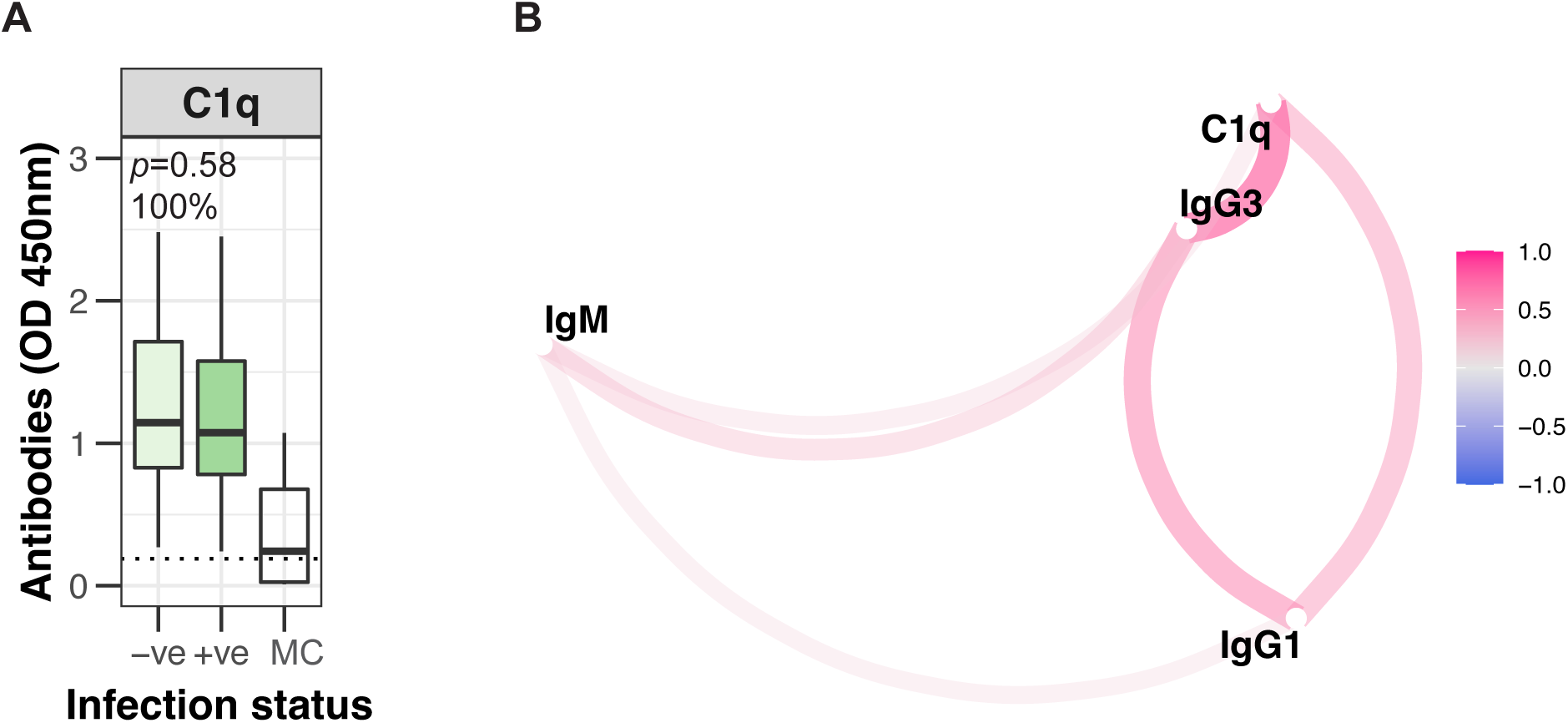
Correlation between acquired antibodies and complement fixation. (**A**) The ability of antibodies to fix complement C1q on the surface of whole gametocytes. Samples were from adults (n=73) residing in Kanyawegi, Kenya and are classified according to infection status (+ve, -ve) for each antibody variable. Malaria-naïve donors from Australia were used as negative controls (MC). C1q levels were measured by ELISA at optical density of 450nm and tested in duplicate; the dotted line represents the antibody positivity threshold (OD levels greater than the upper 95%CI of the mean responses of malaria-naïve Australian controls); % refers to the percentage of samples above the antibody positivity threshold; *p* values comparing antibody groups (infection status) were calculated using an unpaired Mann Whitney test. Box and whisker plots indicate the first and third quartile for the hinges, median line and lowest and highest values no further than 1.5 interquartile range from the hinges for whisker lines. (**B**) In samples from adults residing in Kanyawegi, Kenya, the relationship between antibody variables is visualised using a network plot, which depict the strength and direction of the correlation using Spearman’s rho. The closer the distance between variables, the stronger the correlation. Colour intensity reflects the strength of the correlation coefficients.

### Purified human IgM more strongly mediates complement fixation and activation on gametocytes than IgG

We purified human IgM and IgG from Kenyan adults (n=10) to investigate their respective contributions in mediating complement fixation and activation to gametocytes. IgM and IgG were also purified from malaria naïve Australian donors (n=4) as an experimental negative control. We confirmed the purity of our IgG and IgM fractions compared to commercial human IgM and IgG standards (Supplementary Fig S1A for IgM purification; Fig S1B for IgG purification). The IgM-purified fractions were highly comparable to the commercial human IgM standard and showed little to no bands indicating IgG heavy chain contamination. Similarly, the IgG purified fractions were highly comparable to the commercial human IgG standard and showed little to no IgM contamination. Further, we demonstrated using ELISA with specific IgM detection antibodies that the IgM fractions had limited IgG contamination (Supplementary Fig S1C) and vice versa for the purified IgG fractions (Supplementary Fig S1D).

We confirmed that the purified Kenyan IgM and IgG fractions bound the surface of whole gametocytes (Fig 3A, 3B), which was substantially higher than results using purified IgM or IgG from Australian donors. We measured the ability of the purified antibodies to fix C1q, the first step in the classical complement pathway, and to activate complement leading to formation of the complement complex C5b-C9 (responsible in forming the membrane attack complex to induce cell lysis). Our purified IgM fraction effectively fixed C1q on whole gametocytes (Fig 3C) and promoted formation of the C5b-C9 complex on whole gametocytes (Fig 3D). IgG also promoted fixation and activation of complement. The level of C1q fixation observed was approximately 1.2-fold higher with IgM compared to IgG fractions (OD values were 1.62 for IgM and 1.32 for IgG at 128μg/ml; *p*=0.06). The level of C5b-C9 deposition was 2.7-fold higher with IgM compared to IgG fractions (OD values are 0.73 for IgM and 0.27 for IgG at 128μg/ml; *p*=0.03). Further, relative to the level of C1q fixation, the formation of the C5b-C9 complex was 1.56-fold higher for IgM compared to IgG fractions. Our findings suggest that IgM is more effective than IgG at fixing C1q and promoting C5b-C9 formation.

**Fig 3.**
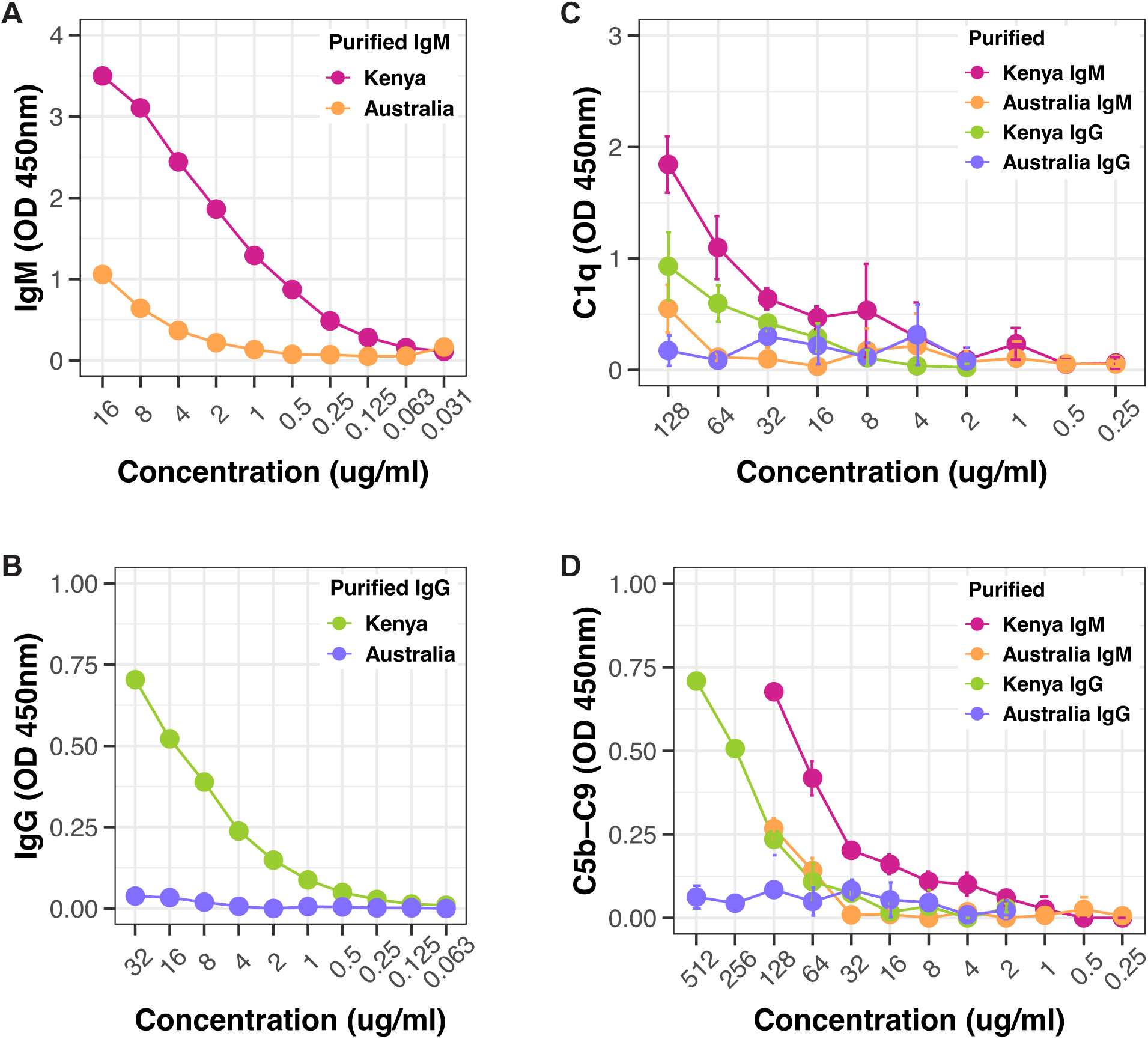
Complement fixation and activation by purified IgM and IgG. IgM and IgG fractions were purified from human sera and tested for reactivity with whole gametocytes. (**A**) IgM purified from individuals from Kenya were highly reactive to whole gametocytes, compared to IgM purified from malaria naïve Australian controls. (**B**) IgG purified from individuals from Kenya were highly reactive to whole gametocytes, compared to IgG purified from malaria naïve Australian controls. Higher levels of (**C**) C1q fixation and (**D**) activation of C5b-C9 were observed with Kenya IgM compared to Kenya IgG. There was minimal reactivity observed with purified fractions of IgM and IgG from Australian controls. IgG, IgM, C1q and C5b-C9 levels were measured by ELISA at optical density of 450nm; (**C**, **D**) data is presented as the mean and standard deviation of samples tested in duplicate. The purified fractions of IgM and IgG were titrated and presented as concentration measured in μg/ml. Assays were performed once to measure total IgM and IgG to whole gametocytes, and twice to measure C1q fixation and C5b-C9 activation with different batches of whole gametocytes (biological replicate).

### IgM is a prominent feature of naturally acquired antibodies to vaccine candidates Pfs230 and Pfs48/45 and other gametocyte surface antigens

To understand the acquisition of naturally acquired antibodies to gametocytes, we assessed the recognition of serum antibodies of current vaccine candidates Pfs230 and Pfs48/45 as well as several other antigens expressed on the surface of gametocytes (Pf38, Pf12 and Pf41). We tested antibody reactivity to 2 recombinant Pfs230 constructs, containing the first 6-cysteine domain of Pfs230^23^ (Pfs230D1) and containing the first two 6-cysteine domains of Pfs230 (Pfs230D1D2). We found that antigen-specific IgG and IgM reactivity among Kenyan adults were very similar between Pfs230D1 and Pfs230D1D2 (Supplementary Fig S2; r_s_=0.73, *p*<0.0001 for IgG and r_s_=0.72, *p*<0.0001 for IgM).

Among adults from Kanyawegi, Kenya (n=102) previously described above, a large proportion were classified as antibody positive for IgG to Pfs230D1 (Fig 4A; 98.4%) and Pfs48/45 (Fig 4B; 98.9%). There was a larger proportion of individuals with IgG3 antibodies compared to IgG1 for both Pfs230D1 (Fig 4A; 84.1% for IgG3 and 66.6% for IgG1) and Pfs48/45 (Fig 4B; 92.1% for IgG3 and 57.9% for IgG1). Further, most individuals were positive for IgM to Pfs230D1 (Fig 4A; 90.5%) and Pfs48/45 (Fig 4B; 87.5%).

**Fig 4.**
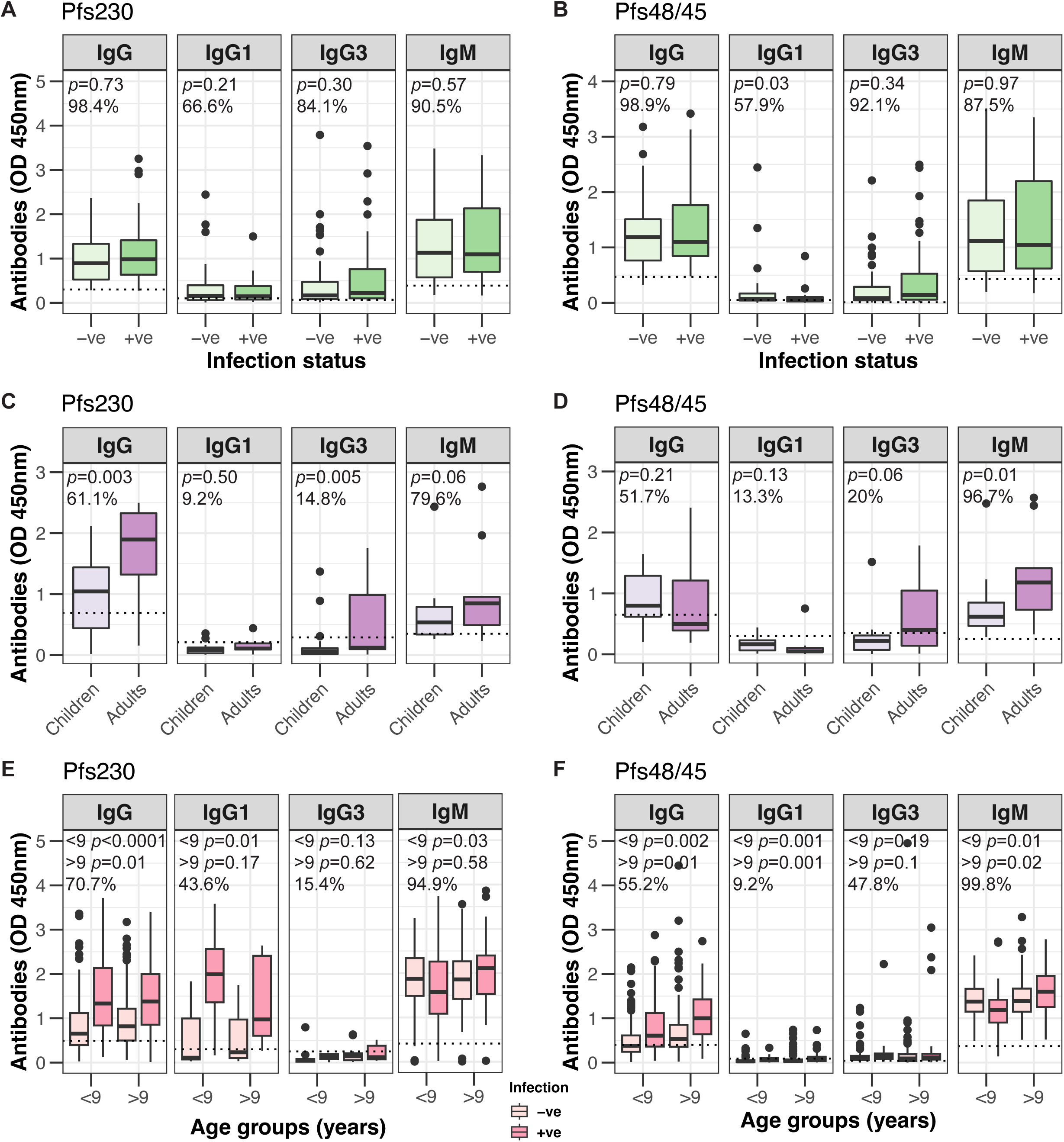
Serum antibodies from Kenya and PNG individuals recognise recombinant *P. falciparum* gametocyte antigens Pfs230 and Pfs48/45. IgG, IgG1 and IgG3 subclasses, and IgM levels were measured to recombinant gametocyte proteins (**A**, **C**, **E**) Pfs230 and (**B**, **D**, **F**) Pfs48/45. Samples were from (**A**, **B**) adults residing in Kanyawegi, Kenya (n=102) and are classified according to infection status (+ve, -ve) for each antibody variable; (**C**, **D**) children (n=49) and adults (n=16) residing in Chulaimbo, Kenya; (**E**, **F**) children residing in Madang, PNG (n=433; n=39 for IgG subclasses to Pfs230; using Pfs230D1D2 construct) and were divided into 2 age groups (<9 years, n=239; >9 years, n=194) and are classified according to infection status (+ve, -ve) for each antibody variable. Antibody levels were measured by ELISA at optical density of 450nm and tested in duplicate; the dotted line represents the antibody positivity threshold (OD levels greater than the upper 95%CI of the mean responses of malaria-naïve Australian controls); % refers to the percentage of samples above the antibody positivity threshold; *p* values comparing antibody groups (infection status in A and B, E and F, or children vs adults in C and D) were calculated using an unpaired Mann Whitney test. Box and whisker plots indicate the first and third quartile for the hinges, median line and lowest and highest values no further than 1.5 interquartile range from the hinges for whisker lines.

In samples from children (n=49) and adults (n=16) from Chulaimbo, Kenya, most individuals were classified as antibody positive for both IgG and IgM to Pfs230D1 (Fig 4C; 61.1% for IgG, 79.6% for IgM) and to Pfs48/45 (Fig 4D; 51.7% for IgG, 96.7% for IgM). There was little IgG1 response to Pfs230D1 (Fig 4C; 9.2%) and Pfs48/45 (Fig 4D; 13.3%). There was a proportion of individuals positive for IgG3 to Pfs230D1 (Fig 4C; 14.8%) and Pfs48/45 (Fig 4C; 20%), and the majority of those responses were in adults. Adults had significantly higher levels of IgG and IgG3 to Pfs230D1 compared to children (Fig 4C; *p*=0.003 for IgG, *p*=0.005 for IgG3), but for Pfs48/45, this was only observed for IgG3 (Fig 4D; *p*=0.06). No difference in IgG levels between children and adults were observed for Pfs48/45 (Fig 4D; *p*=0.21). Adults had higher levels of IgM to Pfs230D1 (Fig 4C; *p*=0.06) and Pfs48/45 (Fig 4D; *p*=0.01) compared to children. When children were divided into age groupings based on their median age, their antibody levels to Pfs230D1 and Pfs48/45 increased with age (Supplementary Fig S3).

We next quantified antibody levels in the PNG cohort using the Pfs230D1D2 protein. A greater proportion of children were classified as positive for IgM to Pfs230 compared to IgG (Fig 4E; 94.9% for IgM vs 70.7% for IgG); similar results were observed with Pfs48/45 (Fig 4F; 99.8% for IgM vs 55.2% for IgG). When classified according to age, older children (>9 years) had significantly higher levels of IgG to Pfs230 (*p*=0.03) and Pfs48/45 (*p*<0.0001), compared to younger children (<9 years). However, this was not observed with IgM levels to Pfs230 (*p*=0.44) or Pfs48/45 (*p*=0.13). Older children also appeared to have significantly higher IgG1 to Pfs230 (*p*=0.005). Further, higher levels of IgG, IgG1 and IgM to Pfs230 (Fig 4E) were observed for younger children (<9 years) who were positive for *P. falciparum* infection, whereas no differences in antibody levels were observed for older children (>9 years). For Pfs48/45 (Fig 4F), higher levels of IgG, IgG1 and IgM were observed for those positive for *P. falciparum* infection in both groups of younger and older children.

We also assessed antibodies to other recombinant 6-cysteine proteins (Pf38, Pf12 and Pf41), that are expressed on the surface of gametocytes. In samples from adults in Kanyawegi, Kenya (n=102), most individuals were classified as antibody positive for IgG (Fig 5A; 89.8%) and IgM (Fig 5A; 88.6%) to Pf38, for IgG (Fig 5B; 88.6%) and IgM (Fig 5B; 89.7%) to Pf12 and for IgG (Fig 5C; 75%) and IgM (Fig 5C; 88.6%) to Pf41. The proportion of individuals classified as antibody positive was higher for IgG3 compared to IgG1 for Pf38 and Pf41 but not Pf12 (Fig 5). There were no significant differences among those classified as positive or negative for *P. falciparum* infection, except for IgG3 to Pf38 (Fig 3A; *p*=0.002).

**Fig 5.**
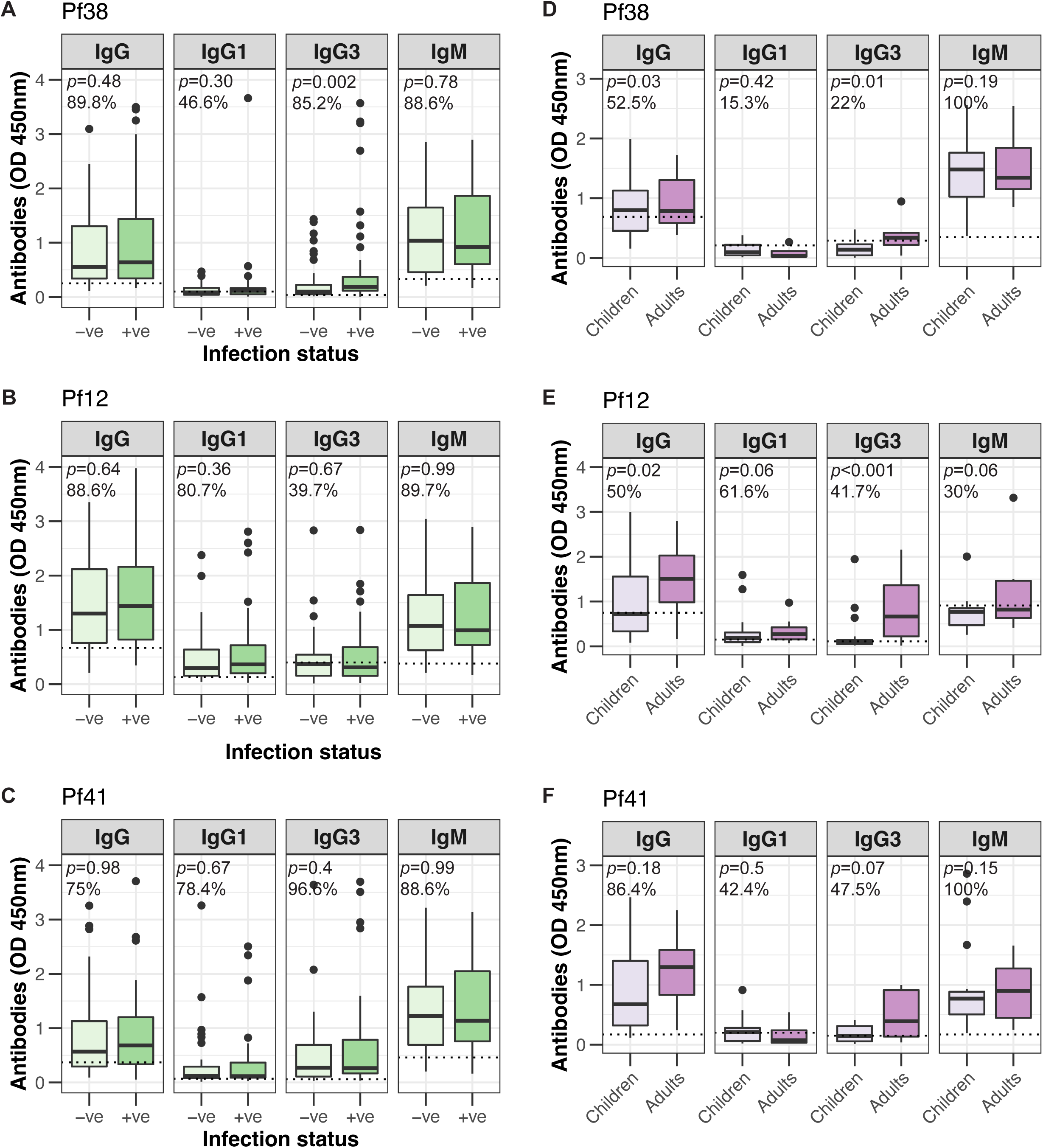
Serum antibodies from Kenya individuals recognise recombinant *P. falciparum* gametocyte antigens Pf38, Pf12 and Pf41. IgG, IgG1 and IgG3 subclasses, and IgM levels were measured to recombinant gametocyte proteins (**A**, **D**) Pf38, (**B**, **E**) Pf12 and (**C**, **F**) Pf41. Samples were from (**A**-**C**) adults residing in Kanyawegi, Kenya (n=102) and are classified according to infection status (+ve, -ve) for each antibody variable; (**D-F**) children (n=49) and adults (n=16) residing in Chulaimbo, Kenya. Antibody levels were measured by ELISA at optical density of 450nm and tested in duplicate; the dotted line represents the antibody positivity threshold (OD levels greater than the upper 95%CI of the mean responses of malaria-naïve Australian controls); % refers to the percentage of samples above the antibody positivity threshold; *p* values comparing antibody groups (infection status or children vs adults) were calculated using an unpaired Mann Whitney test. Box and whisker plots indicate the first and third quartile for the hinges, median line and lowest and highest values no further than 1.5 interquartile range from the hinges for whisker lines.

In samples from children (n=49) and adults (n=16) from Chulaimbo, Kenya, all individuals were classified as antibody positive for IgM directed to Pf38 and Pf41 (Fig 5D, 5F), but only 30% to IgM directed against Pf12 (Fig 5E). The proportion of individuals positive for IgG were lower compared to IgM for Pf38 and Pf41, except Pf12. Adults had significantly higher levels of antibodies compared to children for IgG and IgG3 towards Pf38 (Fig 5D; *p*=0.03, *p*=0.01) and Pf12 (Fig 5E; *p*=0.02, *p*<0.001). Similar patterns of higher antibody responses in adults were observed for IgM to Pf38 (Fig 5D) and IgG, IgG3 and IgM to Pf41 (Fig 5F).

### Strong correlations observed between IgM and complement fixation

We examined the relationship between antigen-specific IgG or IgM with complement fixation in Kenyan adults from Kanyawegi. Network plots were generated for antibody variables to recombinant Pfs230D1 and Pfs48/45 (Fig 6). For responses to Pfs230D1, the correlation between IgM and C1q (r_s_=0.71, *p*<0.0001) was much stronger compared to IgG1 and C1q (r_s_=0.01, *p*=0.94) or IgG3 and C1q (r_s_=0.07, *p*=0.52). Similar results were obtained with responses to Pfs48/45, with IgM and C1q (r_s_=0.48, *p*<0.0001) having a much stronger correlation than for IgG1 and C1q (r_s_=0.20, *p*=0.04) or IgG3 and C1q (r_s_=0.32, *p*=0.001). For both recombinant proteins, weak correlations were observed between antibody variables and age, and negative correlations observed with *P. falciparum* infection status.

**Fig 6.**
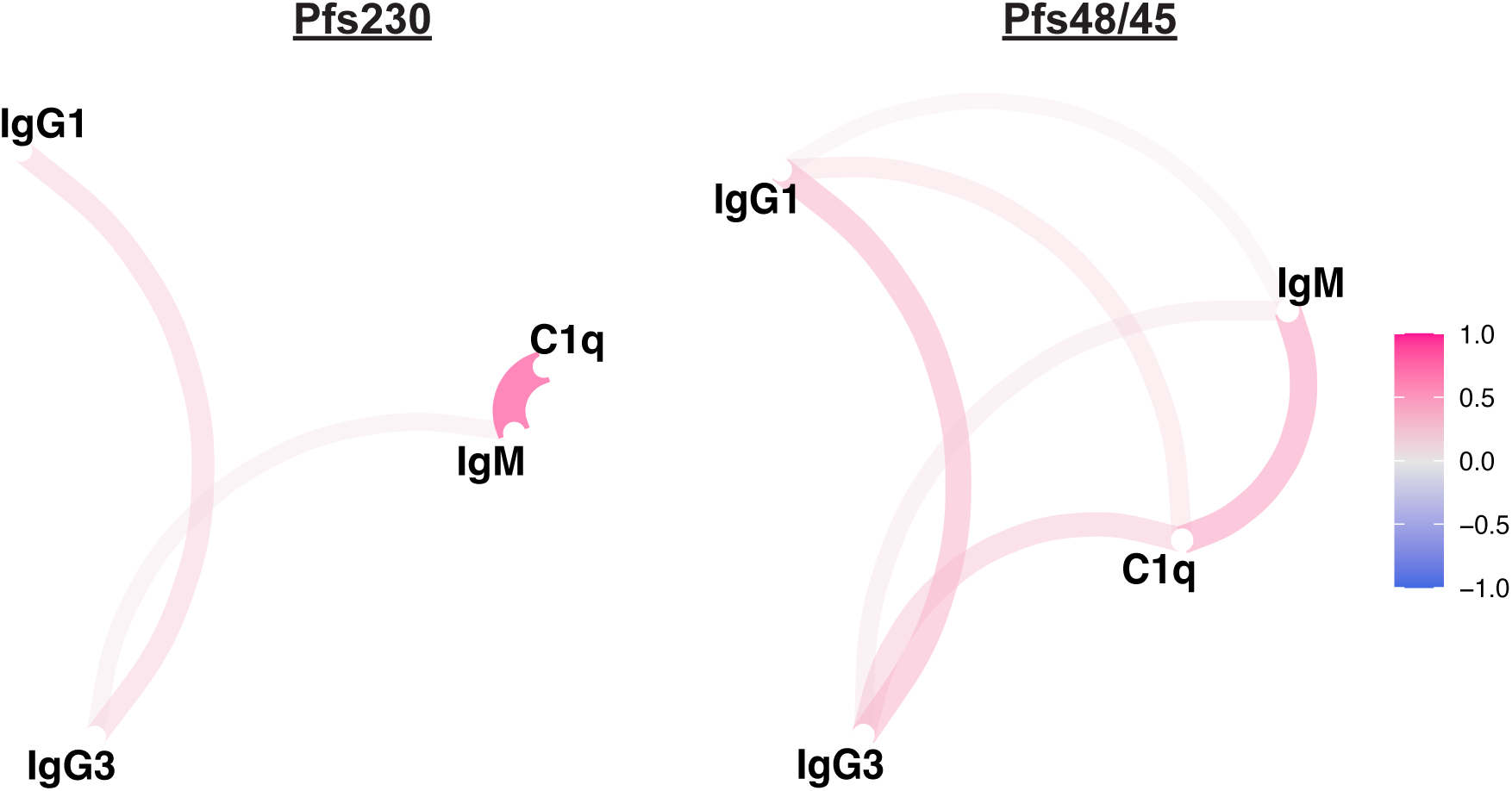
Correlation between antibody variables in Kenyan adults. Network plots depict the relationship between antibody variables; the closer the distance between variables, the higher the correlation (evaluated by Spearman’s rho).

## DISCUSSION

Limited knowledge of the mechanisms and targets of antibodies against *P. falciparum* gametocytes represents a critical roadblock for the advancement of transmission-blocking vaccines. We found that the majority of individuals were capable of mounting an immune response to surface antigens expressed by whole gametocytes, with high levels of IgG, IgG1 and IgG3 subclasses, and IgM responses. The acquired antibody response following natural malaria infections is predominated by cytophilic IgG subclasses IgG1 and IgG3, with little IgG2 and IgG4 observed. Adults and older children generally had higher levels of antibodies, indicating an age-dependent acquisition of immunity in our cohorts. Further, we showed that IgM is a prominent feature of the immune response to gametocytes and recombinant gametocyte proteins. We demonstrated that IgM purified from whole serum was substantially more potent than IgG in fixing complement C1q and promoting activation and the formation of the C5b-C9 complex on gametocytes.

The importance of complement in mediating transmission-reducing activity has been shown by several studies (reviewed in ^6^). A key finding here was that naturally acquired IgM, as well as IgG, could promote complement fixation against gametocytes. We showed that antibodies to whole gametocytes could fix C1q, the first step in the classical complement cascade, and subsequent formation of the C5-C9 complex, which forms the membrane attack complex that can lyse cells. Antibodies and complement in the peripheral blood of the human host are ingested by mosquitoes, and the activation of complement against gametocytes can inhibit further development of mosquito-stage parasites and reduce malaria transmission to other hosts. We also showed that C1q was highly correlated with IgG and IgM magnitude. Comparing the ability of IgG and IgM to activate the complement cascade using purified fractions from malaria-exposed serum, we found that IgM was substantially more potent than IgG at fixing complement C1q and promoting activation and the formation of the C5b-C9 complex. Our findings suggest generating IgM responses may be a potential vaccine strategy for achieving more potent transmission-blocking activity.

We measured the levels of IgG, IgG subclasses and IgM in serum samples to whole gametocytes, and multiple recombinant gametocyte antigens using samples from children and adults from Kenya and PNG. Previous studies have focused on quantifying levels of IgG to Pfs230 and Pfs48/45 only. We showed that human antibodies could recognise the surface of whole, intact gametocytes, with high levels of IgG and IgM, including the cytophilic IgG1 and IgG3 subclasses. Similarly, we found that the majority of samples tested had high levels of IgG and IgM directed at multiple 6-cysteine domain proteins (Pfs230, Pfs48/45, Pf38, Pf12 and Pf41), indicating the acquisition of these antibodies to gametocyte antigens through natural malaria exposure. It was notable that antigen-specific IgM was a prominent feature of the immune response to gametocytes. There was a high prevalence and high levels of IgM to whole gametocytes and recombinant proteins. We previously showed that acquired human IgG could recognise native antigens expressed on the surface of the gametocyte plasma membrane^23^. Naturally acquired antibodies from Gabonese children and adults were also found to recognise the surface of permeabilised erythrocytes infected with mature gametocytes^29^. However, the prominence of IgM in the acquired immune response has not been previously identified. Major gametocyte surface antigens include vaccine candidates Pfs230 and Pfs48/45, which have been previously shown to be targeted by human IgG to mediate transmission-blocking activity quantified using mosquito feeding assays (reviewed in^6,30^).

When individuals from Kenya were classified according to their *P. falciparum* infection status, we did not observe any consistent differences in antibody levels or functional responses to transmission stage antigens. In the PNG cohort of children, higher antibody levels to whole gametocytes were observed for IgG and IgG subclasses, but not IgM, in those classified as infection positive for *P. falciparum* (they typically present with gametocytes as well). However, this was only observed in younger children (<9 years) and not older children (>9 years) suggesting that age plays a key role in antibody acquisition to whole gametocytes. Similarly, younger children (<9 years) who were infection/gametocyte positive had higher levels of IgG, IgG1 and IgM to Pfs230 and Pfs48/45. In older children who were infection/gametocyte positive, higher antibody levels were observed for IgG, IgG1 and IgM to Pfs48/45 but not to Pfs230. Antibodies were generally higher in adults compared to children, indicating an association between antibody acquisition and host age. There are conflicting reports on the association between host age^5^ and the acquisition of antibodies (only IgG reported) to Pfs230 and Pfs48/45^17,31^. Acquired IgG to Pfs230 and Pfs48/45 has been proposed as a measure of recent malaria exposure and increased concurrently with gametocyte density^32^. Future studies in cohorts with samples before, during, and after infection events are needed to better define the relationships between malaria infections and the acquisition of antibodies to gametocytes.

In conclusion, we showed a high prevalence of human antibodies to whole gametocytes and recombinant gametocyte proteins acquired through natural malaria exposure in Kenyan and PNG individuals. Notably, IgM was a prominent feature of acquired immunity to gametocytes, in addition to IgG. Antibodies targeted multiple antigens, including lead vaccine antigens Pfs230 and Pfs48/45, and other members of the 6-cysteine protein family, Pf38, Pf12 and Pf41, suggesting their potential role as key antibody targets in transmission-blocking immunity. We highlighted the importance of IgM responses in mediating functional transmission-blocking immunity, demonstrating that IgM is more potent than IgG in fixing and activating complement against gametocytes, which is an important mechanism in transmission blocking immunity. Serum antibodies to whole gametocytes and recombinant gametocyte proteins were generally higher in adults compared to children, reflecting an age-dependent acquisition of antibodies through natural exposure. Our findings provide substantial new insights to address the major knowledge and innovation gaps required to achieve effective transmission-blocking vaccines for malaria elimination. These findings change the current understanding of transmission-blocking immunity, suggesting an important role of IgM to gametocytes. Developing vaccine strategies that induce functional IgM to gametocytes could be important for achieving potent transmission-blocking immunity.

## MATERIALS AND METHODS

### Study population and ethics statement

Plasma from adults living in the Kanyawegi sub-district, Kisumu Country, Kenya (n=104, age range 18-79; AR) was collected in August 2007, during a period of relatively low malaria transmission^33^. Plasma from children and adults living in the Chulaimbo sub-district, Kisumu County, Kenya (n=75, age range 0.5-5 years for children and 19.6-69.2 years for adults) was collected in February-March 2007^34^. Clinical data on existing *P. falciparum* infections and gametocyte carriage levels were recorded. Plasma was also available from a prospective treatment re-infection study conducted in Madang, PNG, as previously described^25^. This study enrolled children (n=464; age range 4-12 years) from 8 elementary schools. Briefly, children were treated prior to enrolment and plasma samples were collected 3 weeks after enrolment. Clinical data on existing *P. falciparum* infections and gametocyte carriage levels were recorded. Children who are classified as infection positive typically present with gametocytes as well, with infection and gametocyte status being highly correlated. Some samples from these cohorts were not included in this study due to insufficient samples volumes for testing. For assays using only a subset of human samples, the selection of samples was randomly chosen for antibody measurement.

Ethics approval was obtained from the Alfred Hospital Human Research and Ethics Committee, Australia, the Institutional Review Board for Human Investigation at University Hospitals of Cleveland for Case Western Reserve University, USA, the Ethical Review Committee at the Kenya Medical Research Institute and the PNG Institute of Medical Research Institutional Review Board. Written informed consent was obtained from all study participants or their parents or legal guardians.

### *P. falciparum* culture and gametocyte purification

Gametocytes were generated from the NF54 parasite line, according to established protocols^35^. Mature gametocytes (day 13) were harvested for antibody assays and subjected to saponin lysis to permeabilise the parasitophorous vacuole membrane and erythrocyte membrane as previously described^23^.

### Expression of recombinant proteins

Recombinant expression of Pfs230D1 and Pfs230D1D2 was performed in HEK293 Freestyle cells as previously described^23^. Pfs230D1 contains only the first single 6-cysteine unit (covering amino acids S542-G726 of Pfs230)^23,36^, while Pfs230D1D2 contains the first and second 6cysteine unit (covering amino acids S542-T921 of Pfs230). Both constructs contain an N-terminal 6His purification tag and N-linked glycosylated motifs were substituted with serine. Pfs48/45 was expressed in *L. lactis*^37^ and Pf12, Pf38 and Pf41 was expressed in *E. coli*^7^.

### Measuring antibodies to recombinant proteins and whole gametocytes by ELISA

The level of antibodies to recombinant proteins was measured using standard ELISA methods as previously described^23,38^. Briefly, antigens were coated at 0.5 μg/ml in 1 x PBS and incubated overnight at 4°C. Whole gametocytes were coated at 1.2 x 10^6 cells per ml and incubated for 2 hours at 37°C. Plates were blocked with 1% (w/v) casein in 1 x PBS (Sigma-Aldrich) for recombinant proteins and with 1% (w/v) BSA in 1 x PBS for whole gametocytes for 2 hours at 37°C. For recombinant proteins, serum samples were added for 2 hours at room temperature, while for whole gametocytes, serum samples were incubated overnight at 4°C.

For IgG detection, plates were incubated with a goat anti-human IgG HRP-conjugate (1/1000; Thermo Fisher Scientific). For IgG subclasses and IgM detection, plates were incubated with an additional step of mouse anti-human IgG1, IgG3, or IgM (1/1000; Thermo Fisher Scientific) for 1 hour at room temperature, followed by detection with a goat anti-mouse IgG HRP (1/1000; Millipore) for 1 hour at room temperature. Enzymatic activity was detected using TMB liquid substrate (Thermo Scientific) and stopped using 1M sulphuric acid. Antibody levels were measured as optical density at 450nm using the Multiskan Go plate reader (Thermo Fisher Scientific). Washes were performed in between all incubation steps using a microplate washer (Millennium Science, AUS) with 0.05% (v/v) Tween in 1 x PBS, except for assays with whole gametocytes which was washed in 1 x PBS alone. For IgG, IgG subclass and IgM detection, serum samples were used at 1/100 dilution for all recombinant proteins and whole gametocytes. A pool of hyperimmune serum from PNG adult donors was used as a positive control, while serum from malaria naïve Australian adults was used as a negative control.

### Measuring complement fixation and activation by ELISA

The level of complement fixation was measured through established protocols, using purified recombinant C1q^39^. Briefly, antigens were coated at 1 μg/ml in 1 x PBS and incubated overnight at 4°C. Whole gametocytes were coated at 1.2 x 10^6 cells per ml and incubated for 2 hours at 37°C. Blocking and wash steps were performed as described above for standard ELISA. After incubation with human antibody samples (1/50 dilution), purified human C1q (10μg/ml; Millipore) or fresh human serum at 10% (v/v) dilution for C5b-C9 detection were added as a source of complement for 30 minutes at room temperature, followed by a rabbit anti-C1q (1/2000; in-house) or rabbit anti-C5b-C9b (1/1000; (Calbiochem) detection antibody and finally, a goat anti-rabbit IgG HRP (1/2500; Millipore) for 1 hour at room temperature. The level of C1q fixation or C5b-C9 deposition was developed using 3,3’, 5,5’ tetramethylbenzidine (TMB) liquid substrate (Sigma-Aldrich), reactivity was stopped using 1M sulphuric acid and measured as optical density (OD) at 450nm using a Multiskan GO plate reader (Thermo Fisher Scientific).

### Purification of human IgM and IgG fractions

High IgM responders to whole gametocytes from Kenyan adults were chosen and pooled (n=10) for purification. IgM and IgG were also purified from a pool of malaria naïve Australian donors (n=4). Firstly, to purify IgM from whole serum, 100 μl of POROS^TM^ CaptureSelect^TM^ IgM Affinity Matrix packed resin (Thermo Fisher Scientific) was washed thrice with 5 column volume (CV; 500 μl) of 1 x PBS and spinning at 800 x*g* for 5 minutes. 500 μl of diluted Kenyan or Melbourne pooled sample was then incubated with the packed resin for 1 hour at room temperature with end-to-end mixing. Non-IgM material was spun through a Micro Bio-Spin Chromatography Column (Bio-Rad) and washed thrice with 3 CV of 1 x PBS and spinning at <600 x*g* in pulses for 30 seconds. IgM was eluted off the beads by adding 1 CV (100 μl) of 0.1M Glycine (AMRESCO) (pH 2.5), incubating for 15-30 seconds, and spinning at <600 x*g* in pulses for 30 seconds into tubes already containing 10ul of 1M Tris (pH 8.2). Three elution fractions were collected, pooled, and buffer exchanged into 1 x PBS (pH 7.2) using 100kDa Amicon Ultra-15 centrifugal filters (Millipore) and spinning at 4,000 x*g* for 20 minutes at room temperature. Protein concentration of the final purified sample was determined using a NanoDrop One (Thermo Fisher Scientific) using relative absorbance at 280nm with an extinction coefficient of 11.8. The non-IgM material collected as flow-through was used for IgG purification using a Melongel IgG purification kit (Thermo Fisher Scientific) as per manufacturer instructions. Purified IgG samples were similarly quantified using an extinction coefficient of 13.7.

### NuPAGE protein analysis

To assess the purity of the IgM or IgG fractions, the purified samples were run on a gel under reducing conditions. Briefly, samples were prepared to contain 3 μg of purified antibody, 1 μl of 1M dithiothreitol (Thermo Fisher Scientific), 2.5 μl of NuPAGE LDS Sample Buffer (4x) (Invitrogen) and 1 x PBS to make up the volume to 10 μl. Samples were incubated at 95°C for 5 minutes, loaded onto a 4-12% Bis-Tris gel (Invitrogen) and ran at 135 V/200 mA for 1 hour in 1 x MES running buffer (Invitrogen). The gel was then stained with InstantBlue Coomassie (Abcam) for several hours before thoroughly destained with distilled H_2_O overnight and imaged.

### Detecting total IgM and IgG in the purified fractions by ELISA

F96 Maxisorp NUNC-Immuno plates (Thermo Fisher Scientific) were directly coated with the purified IgM or IgG fractions in 1 x PBS and incubated for 3 hours at 37°C. Additionally, the commercial standards Chrompure Human IgG (Jackson ImmunoResearch) and IgM (Jackson ImmunoResearch) were used to create a standard curve for precise sample quantification. The plate was then blocked with 0.1% (w/v) casein in 1 x PBS for 2 hours at 37°C. Isotype-specific antibodies were detected with goat anti-human IgG (1/1000, Invitrogen) or IgM (1/2500, Merck) conjugated to HRP (dilutions performed with 0.1% (w/v) casein in 1 x PBS). The plate was then incubated with 3,3’, 5,5’ tetramethylbenzidine (TMB) liquid substrate (Thermo Fisher Scientific, USA) and shielded from light. The colour-changing reaction was stopped using 1% (v/v) H_2_SO_4_ in distilled H2O (v/v) and OD values were immediately read at 450nm using a Multiskan GO plate reader (Thermo Fisher Scientific). Wash steps were conducted prior to addition of any reagent, using a microplate washer (Millennium Science, AUS) with 0.05% (v/v) Tween in 1 x PBS unless specified otherwise. The Kenyan and Australian purified antibodies were titrated alongside the commercial human IgM and IgG standards of known concentration to create a standard curve from which antibody concentrations of the Kenyan and Australian purified samples could be calculated.

### Measuring purified IgM and IgG fractions against whole gametocytes by ELISA

The level of IgM and IgG to whole gametocytes were measured using standard ELISA methods as mentioned above except for a few modifications. The buffer used for assays with purified IgM and IgG fractions was 1% (w/v) casein in 1 x PBS (Sigma-Aldrich) and IgM was detected using an anti-human IgM antibody directly conjugated to HRP (Jackson ImmunoResearch).

### Statistical analyses

Non-parametric analytical methods were used to evaluate antibody results from human cohort studies as the data generated are not normally distributed. Associations between antibody responses measured were estimated using the Spearman rank correlation coefficient (r_s_). The strength of correlations is defined as weak (0-0.3), moderate (0.31-0.6) or strong (0.61-1). All statistical analyses were performed using Prism version 10 (GraphPad Software Inc) and R Studio. *p* values that compare the difference in OD values for purified IgM and IgG was evaluated using an unpaired Mann Whitney test.

## ACKNOWLEDGEMENTS

We thank all the study participants, their parents and the staff involved in the study from Kenya Medical Research Institute and Papua New Guinea Institute of Medical Research. Blood and human serum used for parasite culture and naïve control sera were provided by the Australian Red Cross Blood Bank (Lifeblood, Melbourne).

