## Supplementary Figures for "IgM plays a prominent role in naturally acquired immunity against *Plasmodium falciparum* gametocytes"

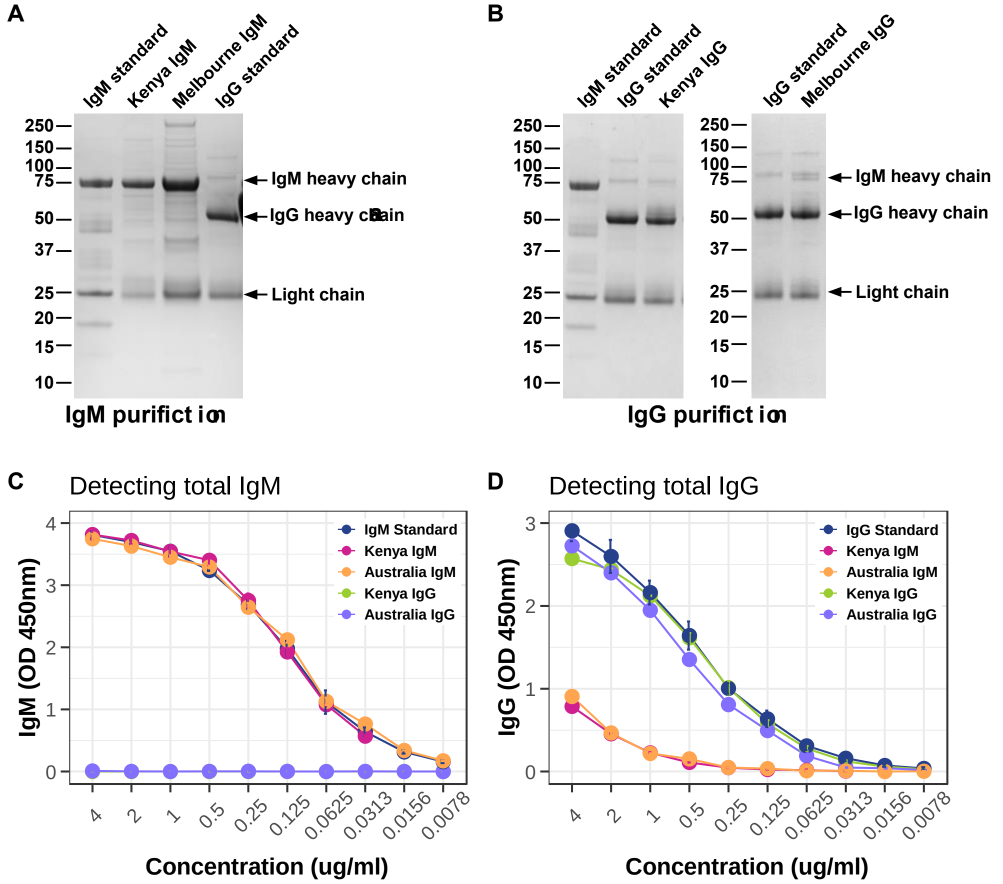


**Fig S1 Validating the purification of IgM and IgG from human sera**

(**A**, **B**) SDS-PAGE to confirm the purify of our human IgM and IgG fractions using commercial standards. Molecular weight markers (kDa) are presented on the left of the gels. Arrows refer to the heavy and light chains of IgM and IgG.

ELISA was used to detect the level of (**C**) IgM and (**D**) IgG purified from pooled Kenya or Australian samples, compared to commercial standards. Purified IgM and IgG fractions were coated directly on the ELISA plate and measured using anti-human IgM or IgG detection antibodies. Antibody levels are measured at optical density (OD) of 450nm; data is presented as mean OD and error bars refer to standard deviation of samples tested in duplicate; concentration (µg/ml) refers to the amount of purified IgM/IgG coated directly onto the ELISA plate.

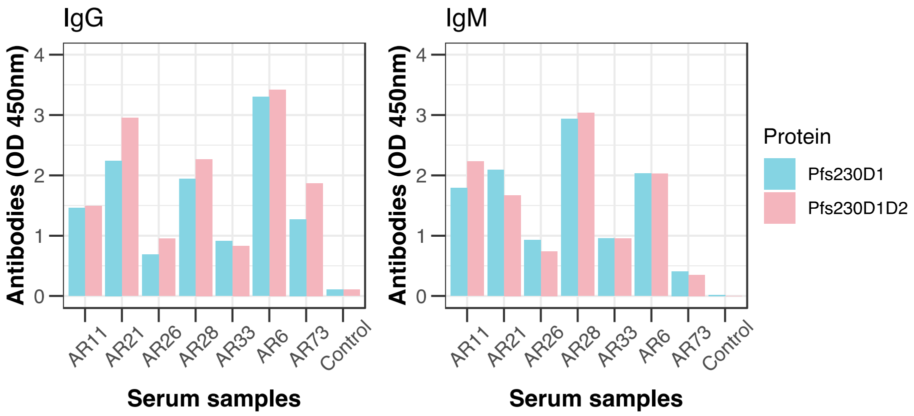


**Fig S2 Comparing antibody levels to 2 different constructs of Pfs230**

IgG (left) and IgM (right) levels were compared between Pfs230D1 and Pfs230D1D2 constructs. Samples were from malaria exposed Kenyan adults (Kanyawegi; AR) and malaria-naive Australian donors (Control); bars represent the mean and range of samples tested in duplicate; antibody levels were measured by ELISA at optical density of 450nm.

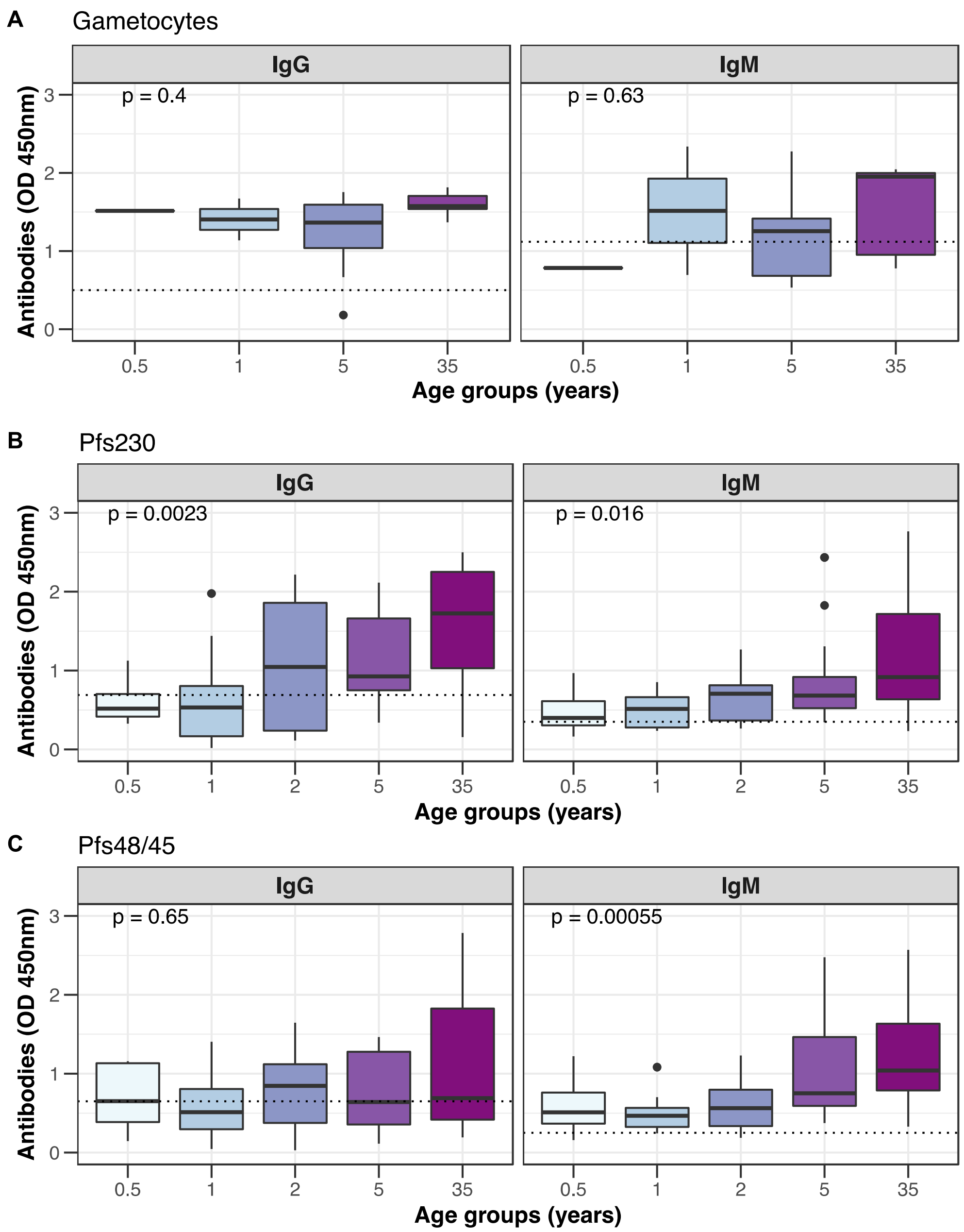


**Fig S3 Serum antibodies from Kenyan children recognise recombinant *P. falciparum* gametocytes and recombinant gametocyte proteins**

IgG and IgM levels were measured to (**A**) whole gametocytes and recombinant gametocyte proteins (**B**) Pfs230D1 and (**C**) Pfs48/45. Samples were from children and adults residing in Chulaimbo, Kenya and were divided into groups based on their median ages of 0.5, 1, 2, 5 and 35 years (n=1, 1, 1, 10, 5 for gametocytes; n=7, 11, 15, 18, 16 for recombinant proteins). Box plots represent the median and interquartile range of samples measured in duplicate; antibody levels were measured by ELISA at optical density of 450nm; the dotted line represents the antibody positivity threshold (OD levels greater than the upper 95%CI of the mean responses of malaria-naïve Australian controls); % refers to the percentage of samples above the antibody positivity threshold; *p* values were calculated using the Kruskal-Wallis test. Box and whisker plots indicate the first and third quartile for the hinges, median line and lowest and highest values no further than 1.5 interquartile range from the hinges for whisker lines.
